# Retrospective Analysis of the Effects of BWF Interdisciplinary Postdoctoral to Faculty Transition Awards on Future Funding Success

**DOI:** 10.1101/2024.07.05.602167

**Authors:** Mandeep K. Sekhon, Melanie Scott, Cynthia L. Green, Miquella C. Rose

## Abstract

Established by the Burroughs Wellcome Fund (BWF) in 2001, the Career Award at the Scientific Interface (CASI) is a career development award for scientists with doctoral training in the physical/mathematical/computational sciences or engineering conducting postdoctoral research in the biological sciences. The goal of the program is to support early career scientists interested in pursuing an independent research career with an interdisciplinary focus. In order to assess the benefit of the CASI award on recipients, the authors undertook a retrospective analysis of the funding data for CASI recipients to evaluate success against matching cohorts. These cohorts included applicants who succeeded to the final interview stage but were ultimately unsuccessful (interviewed), applicants who submitted proposals but did not make it to the final interview stage (proposal declined), and a randomly selected dataset of researchers from a comparable program, the highly competitive Pathway to Independence Award (K99/R00) from the National Institutes of Health (NIH). The results indicate that CASI recipients outperformed unsuccessful applicants and their K99/R00 counterparts in federal grant rates and overall grant dollars. The authors’ conclusion affirms that the CASI mechanism and BWF support successfully achieve the objective of invigorating the careers of young investigators, resulting in tangible downstream long-term effects.

## Introduction

The Burroughs Wellcome Fund (BWF) was an early adopter of the funding model of providing bridge funding to early career researchers. In 1994, BWF launched the Career Awards in the Biomedical Sciences (CABS), to assist postdoctoral scientists with funding during both the postdoctoral phase and into the early independent career as biomedical faculty [1]. Other foundations, such as the Doris Duke Charitable Foundation and the American Heart Association, soon established their own postdoctoral bridging programs. In 2005, the NIH became the first U.S. federal funding agency to provide this type of funding with its Pathway to Independence program [2]. BWF has long had the mission to provide funding in areas of unmet need in the biomedical research space. Since increased funding had become available for biomedical researchers, BWF refocused its resources to the area of interdisciplinary research. Recognizing the value of applying interdisciplinary approaches to address questions in the biomedical sciences, the Career Awards at the Scientific Interface (CASI) program was modeled closely on BWF’s Career Awards in the Biomedical Sciences (CABS) program but was unique in design by the requirement that applicants have doctoral level training or evidence of significant expertise in a computational, theoretical, or physical science discipline outside of biology.

Scientific advances such as genomics, quantitative structural biology, imaging techniques, and modeling of complex systems have created opportunities for exciting research careers at the interface between the physical/computational sciences and the biological sciences. Addressing crucial challenges in biology requires scientists trained in disciplines such as chemistry, physics, applied mathematics, computer science, and engineering. BWF had long recognized this critical area by creating the institutional scientific interface funding mechanism in 1996. This program funded ten academic institutions with a total investment of over $26 million dollars. Each institution had different models for trainees, but all had the purpose to provide interdisciplinary training to young investigators from quantitative, theoretical, and computational backgrounds to introduce innovative ideas and approaches into the biological sciences [3]. After this pilot award mechanism, BWF funded a series of awards for short courses in the area and launched the Career Awards at the Scientific Interface (CASI) in 2002.

In its current version, CASI is a five-year, $560,000 bridging award that funds two years of postdoctoral training and three years of faculty support for early career development of these interdisciplinary researchers. In February 2023, BWF increased the award amount from $500,000 to $560,000 to keep pace with inflation. Additionally, BWF recognized the challenges faced by early-career scientists with caregiving responsibilities and introduced a provision that will allow CASI awardees to allocate up to $5000 per year to cover childcare expenses beginning with the 2023 cohort of CASI awardees. Note that the total award amount during the time of this retrospective analysis was $500,000.

BWF was interested in investigating the effect of the Burroughs Wellcome Fund CASI award on awarded investigators’ independent careers and contributions to science. The ability to secure federal funding compared to similar cohorts of researchers considered was analyzed using several funding metrics, such as total grant number, dollars, and annual yearly grant rate.

## Materials and Methods

### Data collection

Publication and funding data were obtained from the Digital Science Dimensions Research Information System [4]. Briefly, BWF provided a list of CASI program applications with 513 unique applicants from 2007-2017. These applicants were broken down into the following categories: awardees (n=109; 21.2%), interviewed and declined (n=117; 22.8%) and proposal declined (n=287; 55.9%). Duplicates within each category (for example, proposal declined more than once) were removed. Digital Science matched a comparable number (n=125) of NIH Pathway to Independence Award K99/R00 from the same time period (2007-2017) for a total of 638 unique investigators. K99/R00 awardees were randomly selected from the K99/R00 National Institute of General Medical Sciences (NIGMS) applicant pool to control variables such as funding line differences between NIH Institutes and Centers (ICs). Specifically, Digital Science used PostgreSQL to generate a random number associated to each of the NIGMS K99/R00 award recipients with award start years listed above, then sorted numerically by the randomly assigned number and selected the top 125 records. The year estimates the approximate award appointment year based on year of BWF application or K99/R00 award. The 638 investigators were then analyzed by Digital Science to obtain researcher data. To study years without the NIH comparison, CASI applications were provided from 2002-2017. From these applications, we have data from a total of 700 applicants, 145 (20.7%) from the awarded group, 156 (22.3%) interviewed/declined, and 399 (57.0%) from the proposal declined group.

Digital Science matched applicants to researcher profiles in Dimensions using a fully automated, rule-based approach. The program partially disambiguates researchers associated with publications and/or grants, and errs on the side of higher precision, lower recall for the person disambiguation available in the user interface [5]. The resulting data matches were binned from 0-7, with 1 being most confident of a data match and 7 being least confident. In order to ensure the most robust data, match type bins 6 and 7 were discarded and not used in further analysis. Data was categorized for the K99/R00 recipients in the data bin 0, since the data were a direct download from NIH and there was no ambiguity as to their identity being a match.

### Statistical analysis

Categorical data are presented as counts with percentages, while continuous data are presented using the mean, standard deviation (SD), median, 25^th^-75^th^ percentiles (Q1-Q3) and range. The association of receiving a federal grant (yes vs. no) and CASI awardee cohort was compared using a chi-square test. Generalized linear regression models with either a Poisson (annual grant rate) or negative binomial (annual funding) distribution were used to compare CASI awardee cohorts after adjusting for the estimated appointment year. Results are presented as the least square mean (LSM) estimate with 95% confidence interval (CI) for each cohort. A post-hoc simulated alpha adjustment was used to control the type I error for multiple pairwise comparisons between cohorts. Since the regression models are based on a logarithmic distribution, the difference between groups is given by the percentage increase or decrease in the expected mean estimate with 95% CI. Statistical analyses were done using SAS version 9.4 (SAS Institute, Inc., Cary, NC), and a p-value <0.05 was considered statistically significant.

## Results

The Career Awards at the Scientific Interface is a grant mechanism intended to support the early career development of researchers interested in scientific questions of an interdisciplinary nature. Candidates draw from their training in a scientific field other than biology to propose innovative approaches to answer important questions in the biological sciences. The advisory committee is composed of a multidisciplinary panel of accomplished scientists. The grant mechanism is a two-stage process, in which applicants submit a brief pre-proposal, and then a select number are invited to submit a full proposal. BWF usually receives around ∼250 pre-proposals each cycle, of which approximately ∼30% are invited to submit full proposals.

From this group, a smaller group is further selected (∼20 applicants per year) to present their proposal to an advisory committee (Fig 1a), and approximately 10-11 finalists are chosen to receive an award each year. Applicants for this analysis were chosen from our database from all applicants who submitted proposals from 2002-2017 but were limited to 2007-2017 when comparing to NIH K99/R00 awardees from the same period. Applicants were chosen from the following categories: Proposal Declined, Interviewed/Declined, or Awarded. Proposal Declined (PD) describes any candidate that was declined from the full proposal stage.

**Fig 1.**
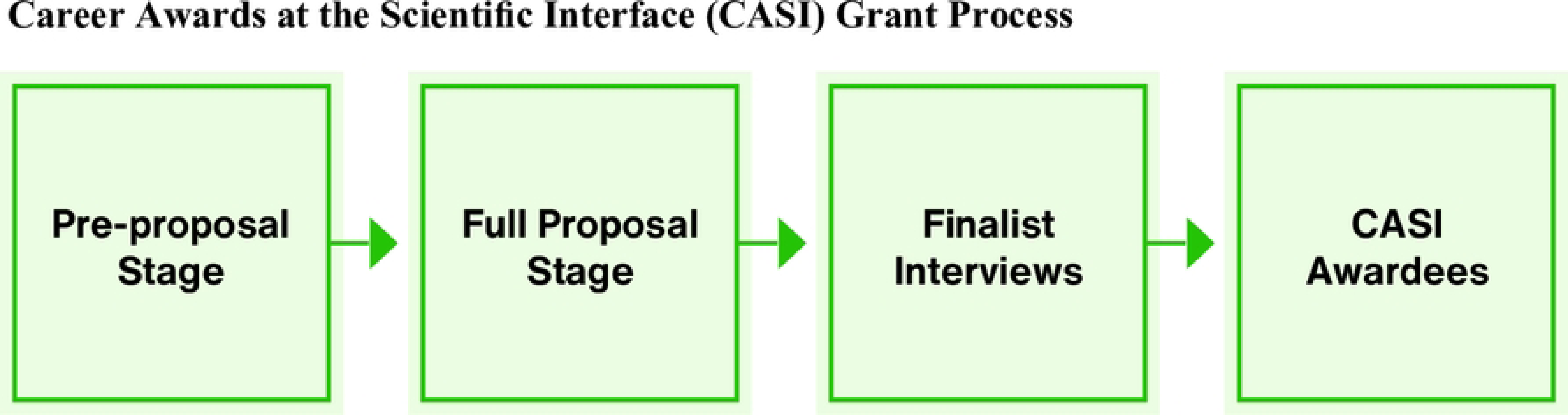
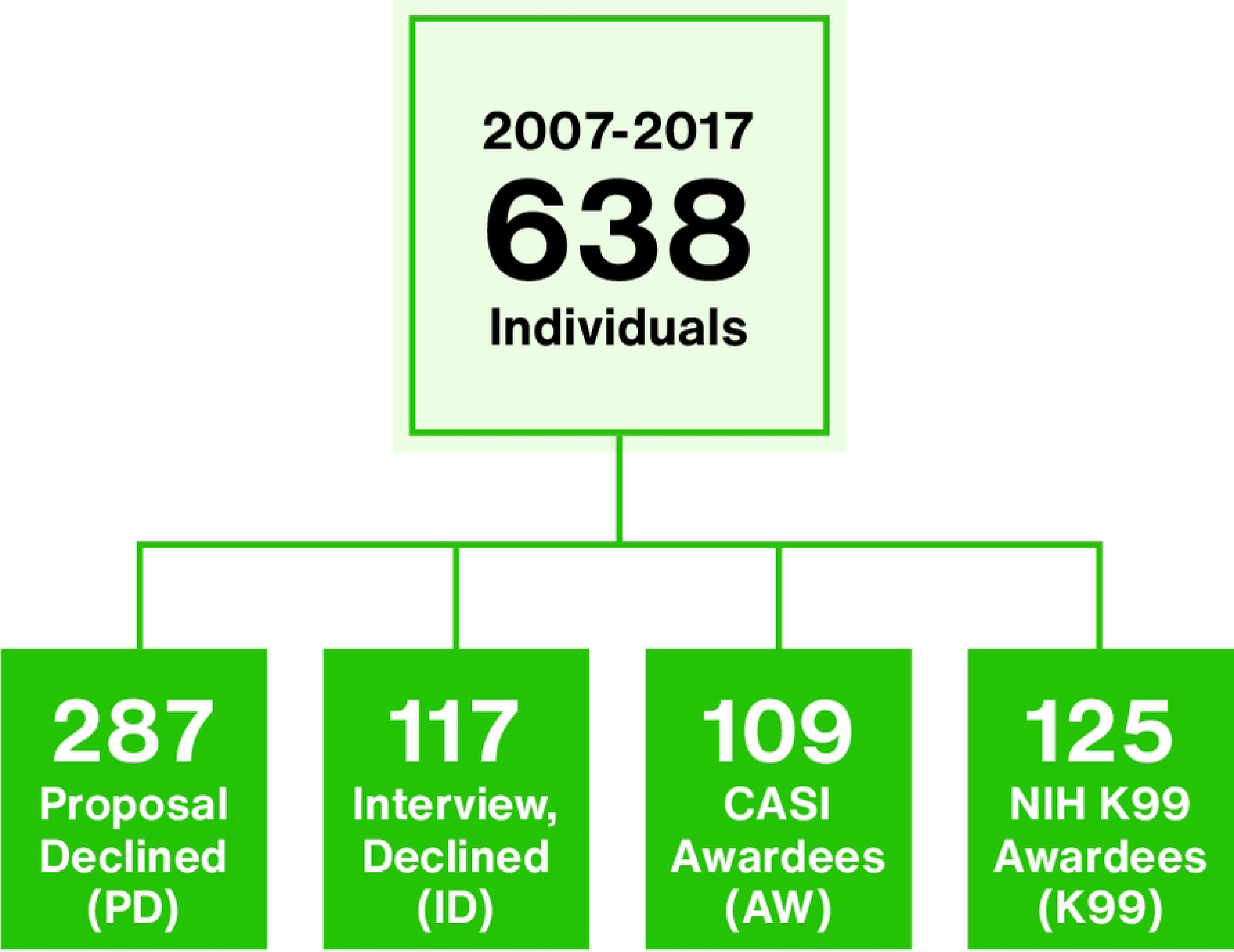
Overview of the Career Awards at the Scientific Interface (CASI) Process, 2007-2017. 1a. Schematic of the Career Awards at the Scientific Interface two stage grant process. 1b. A total of 638 investigators from 2007-2017 were analyzed in this study and the figure shows the counts by cohort.

Interviewed/Declined (ID) describes highly qualified candidates that presented their proposal before the advisory committee but were not chosen for an award, usually due to funding limitations. Awarded (AW) describes candidates that went through all stages and were chosen for the award (Fig 1b). Additionally, we obtained data from a comparable outside funding mechanism, the NIH Pathway to Independence Award K99/R00 during the same funding years (2007-2017). A breakdown of the investigator distribution by category and year can be found as Supporting Information Table 1. A total of 638 individuals were included in further analyses.

**Table 1.**
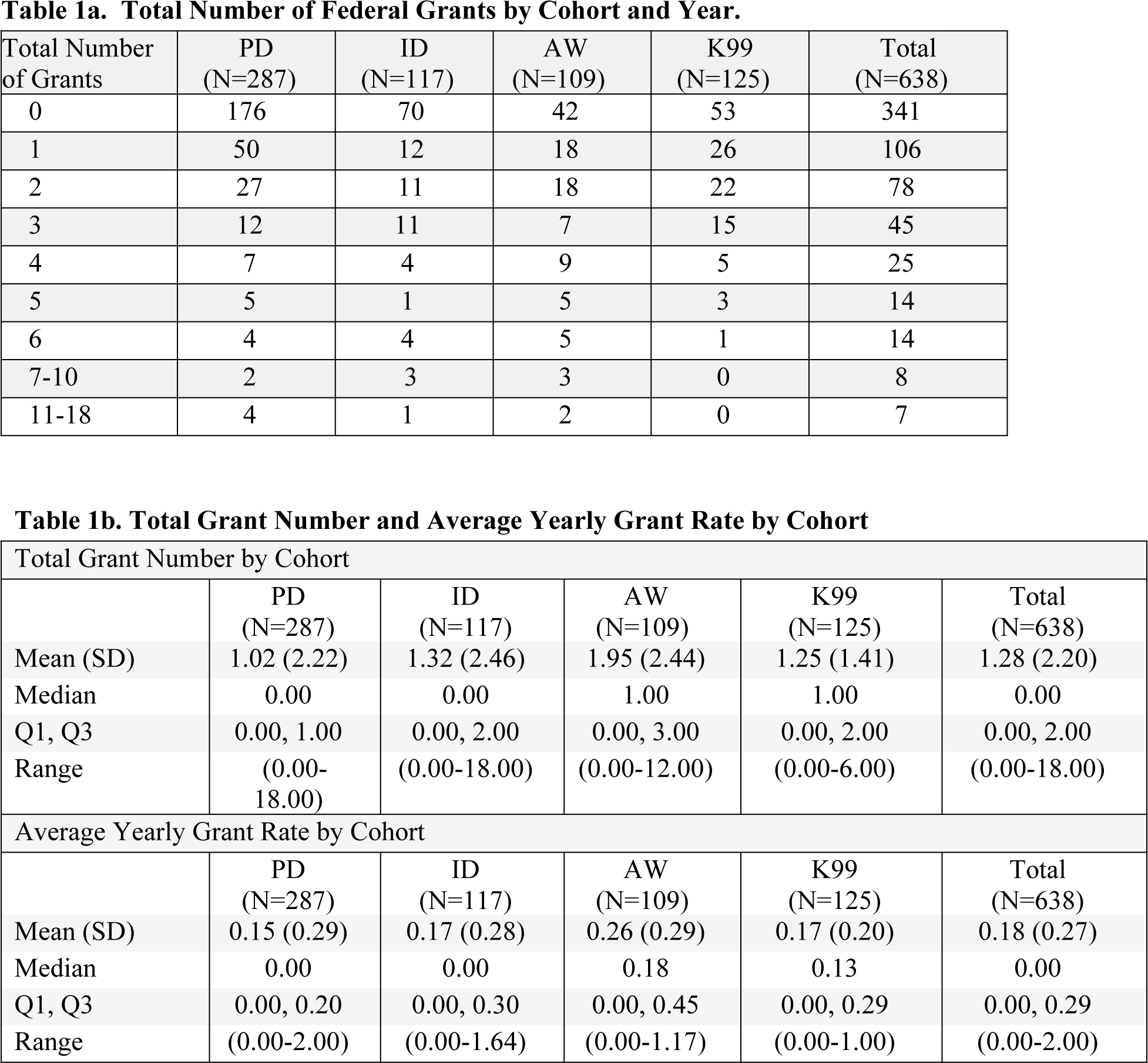
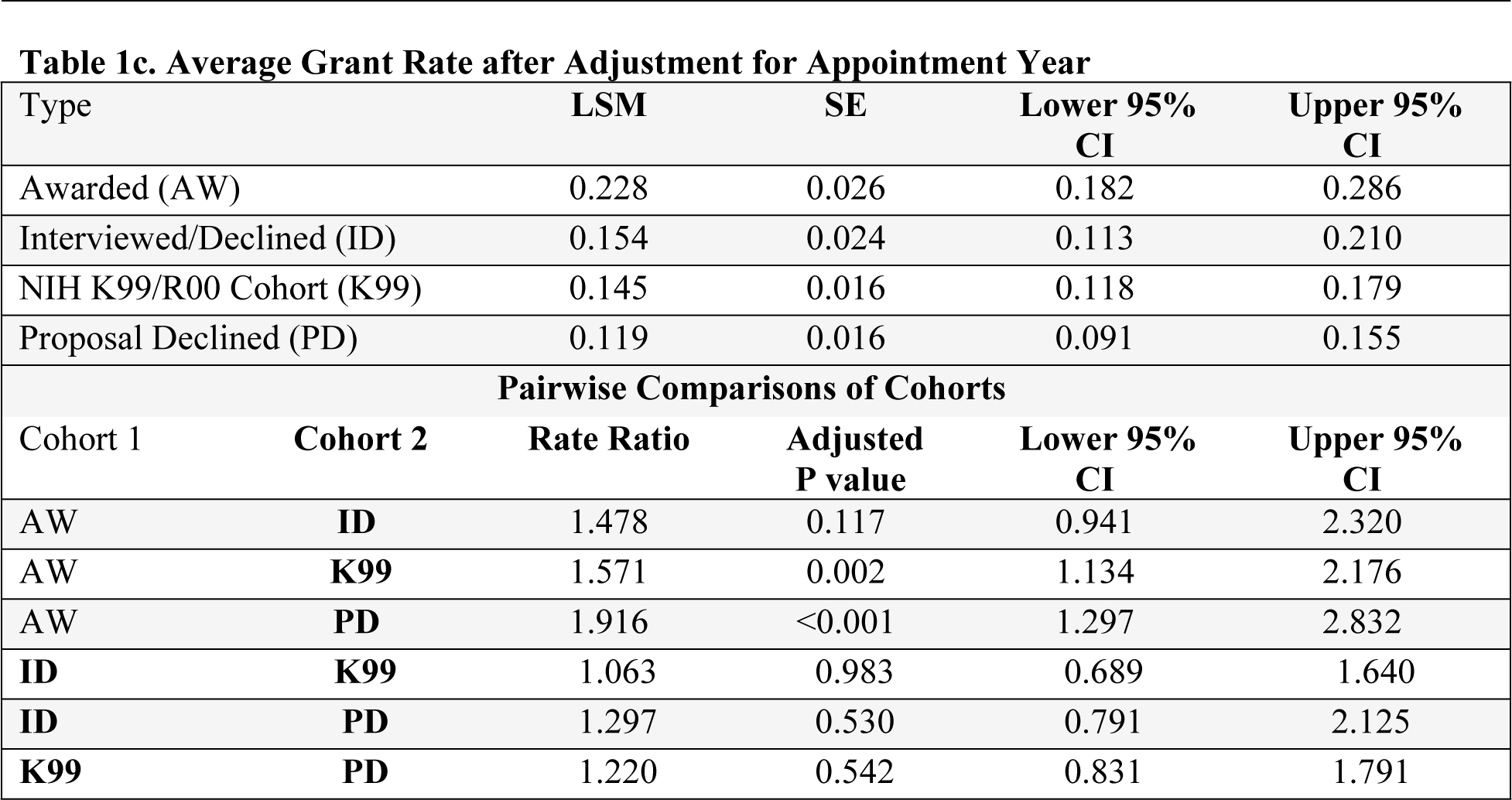
Total Numbers of Grant and Grant Rates. The total and average number of grants for each of the 638 applicants identified by cohort are detailed in the following tables. 1a. Total number of federal grants held by investigators separated by cohort. 1b. Total mean, median, quartiles and ranges of both total grant number and average yearly grant rate separated per cohort. 1c. A generalized linear regression model with Poisson distribution was used to analyze the average grant rate after adjusting for the estimated appointment year. The least square mean (LSM) estimates with 95% confidence intervals (CI) are given for each group (top table) as well as the pairwise differences (rate ratio) between groups (bottom table). AW=Awarded, PD=Proposal Declined, ID=Interviewed/Declined and K99= NIH K99/R00 cohort.

Our first aim was to investigate the number of federal grants that the applicants were awarded throughout their career. The NIH K99/R00 awardee applicants had 1-2 grants entered for the original K99/R00 grant, which could overestimate the amount of federal funding received. Also, many applicants receive federal awards for graduate study, which would not be relevant when analyzing the effect of postdoctoral funding on future federal awards. In order to accurately estimate follow-up federal funding, we eliminated any grants occurring prior to the estimated “appointment” year of the R00, or faculty portion of the award. The total grant number for individuals within the different cohorts is shown in Table 1a. For 638 applicants, 815 grants were identified within the match type process described in the methods section. Total grants held by individuals ranged from 0-18, with over half (341/638; 53.4%) of investigators having zero federal grants. We next looked at the total number of grants and yearly grant rate. The mean ± SD number of grants ranged from 1.02 ± 2.22 for the PD group, 1.32 ± 2.46 for the ID group, 1.95 ± 2.44 for the AW group, and 1.25 ± 1.41 for the K99/R00 group (Table 1b). When the average yearly grant rate was estimated, the mean average grant rate was 0.15 ± 0.29 for the PD group, 0.17 ± 0.28 for the ID group, 0.26 ± 0.29 for the AW group, and 0.17 ± 0.20 for the K99/R00 group (Table 1b). Overall, the CASI AW group received more grants at a higher annual rate than the other groups on average. In order to eliminate the chance that some extremely high achieving senior academics in older cohorts were skewing the data, we ran an analysis to compare each cohort while controlling for the estimated appointment year using a generalized linear regression model (Table 1c). The AW cohort had a significantly higher expected annual rate (LSM 0.23; 95% CI: 0.18-0.29) when compared to both the PD (LSM 0.12; 95% CI: 0.09-0.16) and K99/R00 (LSM 0.15; 95% CI: 0.12-0.18) groups. Though not statistically significant, the expected rate was also higher in the AW than the ID (LSM 0.15; 0.11-0.21) group. In this analysis, the CASI awardee (AW) expected mean grant rate is 1.57 (95% CI: 1.13-2.18; p=0.002) times higher than NIH K99/R00 awardees, and 1.92 (95% CI: 1.30-2.83; p<0.001) times higher than proposal declined (PD) group. The AW group mean rate was 1.48 (95% CI: 0.94-2.32) times higher than the interviewed declined group, but this was not statistically significant (p=0.117). There were no other notable differences between groups that even remotely approached statistical significance (data not shown).

When investigating the total funding dollars, some interesting trends appear. The median total funding dollars was $0 for the proposal declined (PD) group, as well as for the interviewed but declined (ID) group. The median funding for the CASI awarded group (AW) was $689,117, and the median for the K99/R00 group was similar at $636,300 (Table 2a). When comparing the third quartiles across groups, the proposal declined group was funded at $592,520, the interviewed but declined were slightly higher at $883,880, the K99/R00 was at $2,040,487, with the CASI awarded group having the highest value at the third quartile with $2,763,931. Average yearly funding had a similar trajectory, with both the median funding for proposal declined and interviewed declined at $0, and the CASI Awarded and K99/R00 awardees with similar annual federal funding rates ($92,171 and $91,154, respectively). These combined data are likely due to the funding payout of a typical 5-year NIH R01 or R01 equivalent grant, which was $559,680 in 2020, according to a 2021 analysis [6].

**Table 2.**
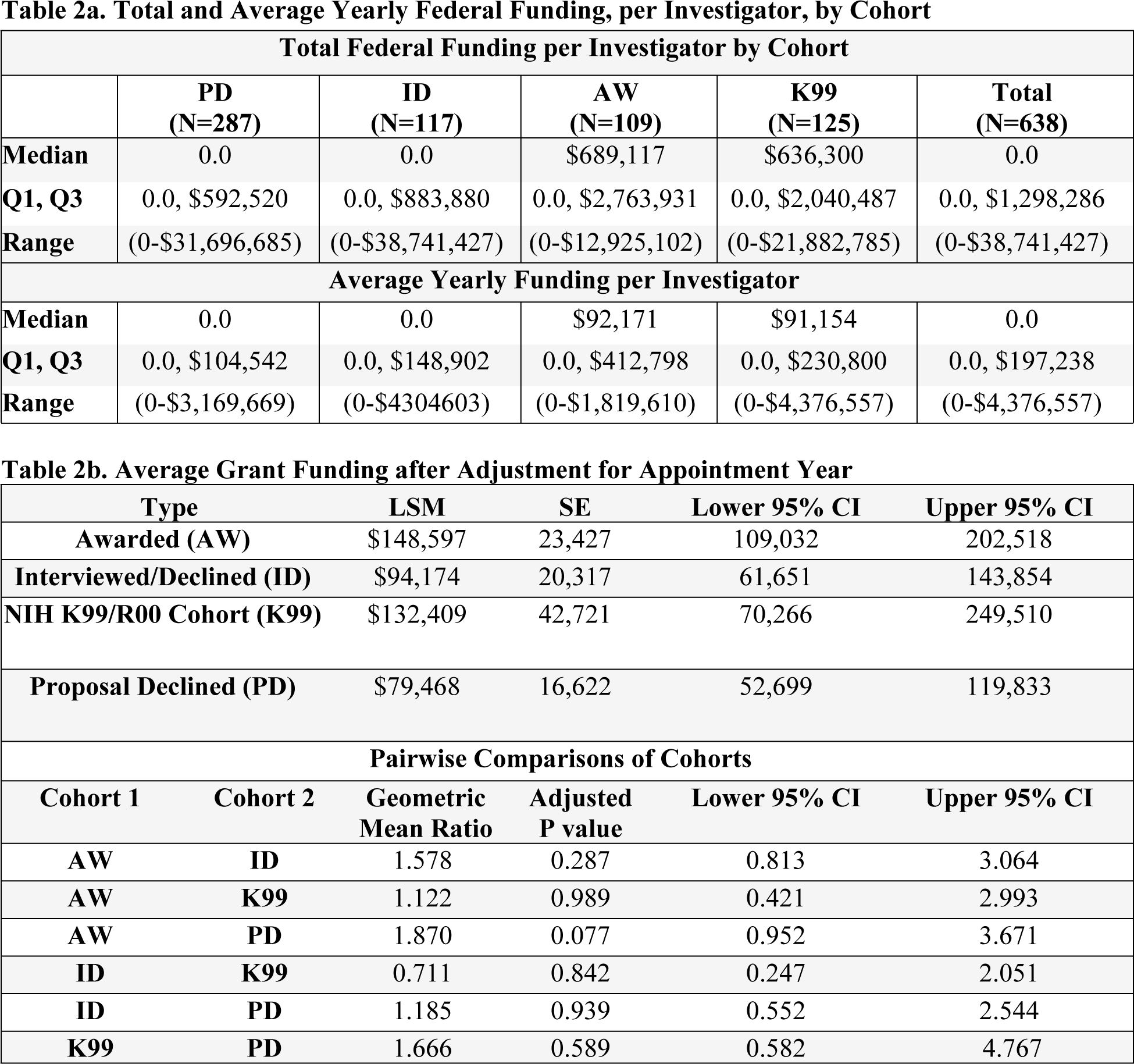
Grant Funding Dollar Amounts 2a. Total median, quartiles and ranges of both total grant funding and average yearly grant dollars were separated per cohort. 2b. A generalized linear regression model with negative binomial distribution was used to analyze the average grant rate after adjusting for the estimated appointment year. The least square mean (LSM) estimates with 95% confidence intervals (CI) are given for each group (top table) as well as the pairwise differences (geometric mean ratio) between groups (bottom table). AW=Awarded, PD=Proposal Declined, ID=Interviewed/Declined and K99= NIH K99/R00 cohort.

A generalized linear regression model was used to analyze the average annual grant funding amount after adjusting for the estimated appointment year (Table 2b). In this analysis, CASI awardees averaged $148,597 dollars/year (95% CI: $109,032-202,518), compared to $132,409 for NIH K99 awardees (95% CI: $70,266-249,510), $94,174/year (95% CI: $61,651–143,854) for the interviewed and declined, and $79,468 (95% CI: $52,699-119,833) for the proposal declined group. After adjusting for the estimated appointment year, as was done in the previous analysis, none of the groups were found to be statistically significant in pairwise testing (Table 2b, bottom panel). However, the trends were similar to the previous grant rate results with the AW group having 1.87 times higher expected annual funding (95% CI: 0.95-3.67; p=0.077) compared to the PD group.

One conclusion could be that when comparing cohorts against each other, they are remarkably similar with only minor statistically significant differences in total funding yet no statistical difference in yearly funding. Another conclusion could be that the yearly funding is not statistically significant because the awardees are still too early in their career to see meaningful differences. While we cannot compare the data with K99/R00 applicants any earlier than 2007 because that is the first year it was launched, we can go back to data from as early as 2002 for all BWF applicants. From these applications, we have data from a total of 700 applicants, 145 (20.7%) from the awarded group, 156 (22.3%) interviewed/declined, and 399 (57.0%) from the proposal declined group (Figure 2). The breakdown by group type and year can be found in Supplemental Digital Content Appendix 2. In this group of 700, a total of 291 (41.6%) applicants did not have federal funding (Table 3a). When we break this down by cohort, only 35 (24.1%) CASI awardees did not have federal funding, as compared to 65 (41.7%) from the interviewed and declined group and 191 (47.9%) from the proposal declined group. The majority of applicants (n=23/35; 65.7%) without federal funding in the CASI awarded group (AW) were newer investigators from 2015-2017. The distribution in the interviewed and declined group (ID) and proposal declined group (PD) were more evenly spread amongst cohorts from 2002-2017.

**Figure 2.**
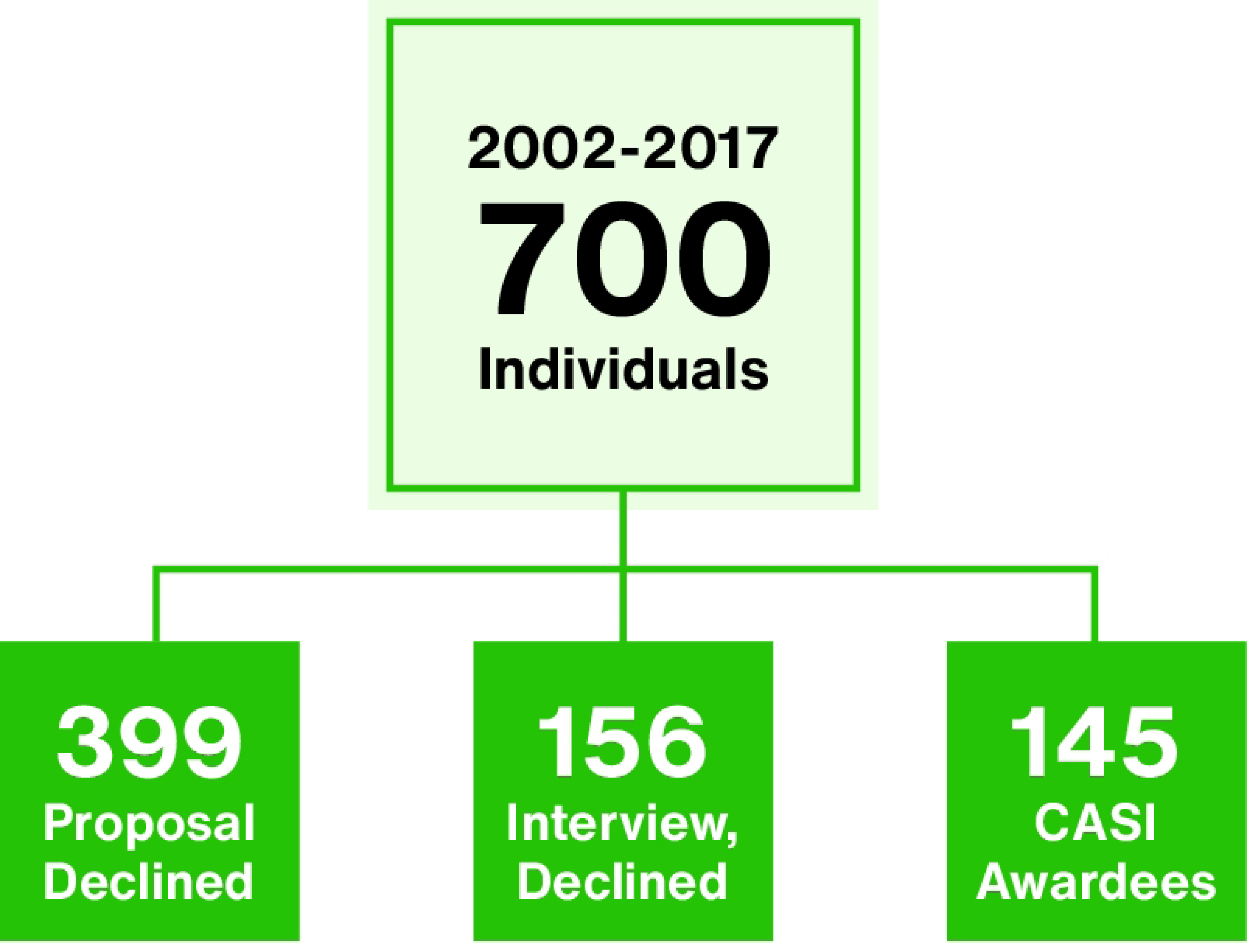
CASI Grant Applicants from 2002-2017. Schematic of the Career Awards at the Scientific Interface two stage grant process. A total of 700 investigators from 2002-2017 were analyzed in this study and the figure shows the counts by cohort.

**Table 3a.** Funding Success for CASI Grant Applicants from 2002-2017. Number of investigators without federal grants broken down by cohort status and award year. Table 3b. Pairwise Comparison by Cohorts for both Grant Rate and by Grant Dollars for BWF applicants from 2002-2017.

|  | **AW**<br>**(N=35)** | **ID**<br>**(N=65)** | **PD**<br>**(N=191)** | **Total**<br>**(N=291)** |
| --- | --- | --- | --- | --- |
| **Year** |  |  |  |  |
| **2002** | 0 | 2 | 11 | 13 |
| **2003** | 2 | 4 | 11 | 17 |
| **2005** | 3 | 3 | 11 | 17 |
| **2006** | 0 | 1 | 12 | 13 |
| **2007** | 0 | 3 | 10 | 13 |
| **2008** | 0 | 1 | 14 | 15 |
| **2009** | 0 | 4 | 16 | 20 |
| **2011** | 1 | 6 | 8 | 15 |
| **2012** | 1 | 3 | 12 | 16 |
| **2013** | 2 | 5 | 13 | 20 |
| **2014** | 3 | 5 | 17 | 25 |
| **2015** | 8 | 9 | 19 | 36 |
| **2016** | 7 | 11 | 20 | 38 |
| **2017** | 8 | 8 | 17 | 33 |

| **Cohort 1** | **Cohort 2** | **Rate Ratio** | **Adjusted P value** | **Lower 95% CI** | **Upper 95% CI** |
| --- | --- | --- | --- | --- | --- |
| **AW** | **ID** | 1.683 | 0.002 | 1.152 | 2.459 |
| **AW** | **PD** | 2.019 | <0.001 | 1.482 | 2.752 |
| **ID** | **PD** | 1.200 | 0.622 | 0.815 | 1.766 |
| **Grant Dollars Pairwise Comparison by Cohort 2002-2017** |  |  |  |  |  |
| **Cohort 1** | **Cohort 2** | **Geometric Mean Ratio** | **Adjusted P value** | **Lower 95% CI** | **Upper 95% CI** |
| **AW** | **ID** | 2.155 | 0.005 | 1.216 | 3.818 |
| **AW** | **PD** | 2.170 | 0.001 | 1.269 | 3.713 |
| **ID** | **PD** | 1.007 | >0.99 | 0.540 | 1.880 |

When we compare each cohort against their counterparts, we see a clear advantage from receiving an award after adjusting for the appointment year. The CASI awarded group grant rate was 1.68 (95% CI: 1.15-2.46; p=0.002) times than the interviewed and declined group and 2.02 (95% CI: 1.48-2.75; p<0.001) times higher than the proposal declined group (Table 3b). The expecting annual funding dollar amount was also statistically significantly higher, with the CASI awarded group 2.16 (95% CI: 1.22-3.82; p=0.005) times higher than the interviewed and declined group and 2.17 (95% CI: 1.27-3.71; p=0.001) times higher than the proposal declined group (Table 3b).

## Discussion

Analysis of the CASI awardee cohorts against the NIH K99/R00 grant cohort indicates a clear association of the CASI award with success of federal funding with respect to both the funding rate and annual total dollar amount. Although in the early years of the award, only a slight advantage can be seen between CASI recipients and their counterparts (Table 2b), by the time these investigators are more senior in their careers, the differences are quite apparent, with CASI recipients outperforming their counterparts in grant dollars (Table 3b). One explanation for this could be that the CASI award funding gives early career scientists the flexibility to take more risks in their scientific inquiries. The ability to generate preliminary data as a result catalyzes future funding opportunities. Self-reported overall career satisfaction by CASI recipients is likely a result of the positive net effect of the award on career successes and trajectories.

Several groups including BWF have stressed the importance and value of early career support for biomedical scientists [7,8]. Recent literature has demonstrated in the biomedical sciences the relationship between early career awards and increased success in obtaining further funding [9–11]. For basic biomedical scientists, studies have suggested early career awards leading to decreased time from postdoctoral stage to first faculty position [12], perceived ability to obtain faculty jobs [1] and overall perceived career satisfaction [13]. Researchers eligible for the CASI program have additional unique challenges, including working across disciplines, which can lead to lower funding rates [14]. An additional hurdle for interdisciplinary scientists is the trend that there has been a large decline in the interest of pursuing academic careers in general, and a larger percentage of decline for researchers in non-life sciences in particular [15]. We found that the interdisciplinary nature of the CASI recipients did not hinder them compared to the K99/R00 awardees, who as NIH awardees tend to more traditionally focused on basic biomedical research and would be expected to have greater funding than interdisciplinary researchers. In actuality, the CASI recipients grant rate was 1.6 times higher than K99/R00 counterparts (Table 1c), and their median research dollars were not statistically different from their K99/R00 counterparts ($92,171 vs. $91,154, Table 2a). One rationale could be that the availability of this guaranteed flexible faculty funds from the CASI bridging award could make these postdoctoral fellows able to obtain preliminary data on higher risk yet highly fundable projects. Another possibility is that securing a prestigious private foundation grant could serve as a “halo effect” and increase chances for follow up funding [16]. Indeed, when we compare the more senior cohorts from 2002-2017 (excluding the K99 cohort), we see a statistically significant difference between the awarded group and the interviewed and declined. It is important to note that these two groups both represent the top interdisciplinary postdoctoral applicants each year, and the talent pool is often much larger than the amount allotted budget. However, with that in mind, over time, the awarded group’s grant rate was 1.68 (95% CI: 1.15-2.46) that of the interviewed and declined and the annual funding rate was awarded group 2.16 (95% CI: 1.22-3.82; p=0.005) times higher than the interviewed and declined group. While beyond the scope of this study, an interesting future direction would be to compare types of follow-up funding, to determine if previous funding from philanthropy increases the likelihood of successful funding from other private and public philanthropic groups. Overall, the goal to advance early career researchers working at the intersection of physical, mathematical, computational and/or engineering and the biological sciences are clearly achieved through the CASI award mechanisms.

## Conclusions (optional)

Our findings indicate that the CASI award is meeting the goals of serving as a pathway for early career scientists to launch their careers and serve as a bridge between postdoctoral and early faculty work. To date, approximately 206 CASI awards have been made signifying the commitment of the Burroughs Wellcome Fund to support early career scientists in areas working at the intersection of traditionally disparate fields.

## Acknowledgements

The authors acknowledge former Burroughs Wellcome Fund President John Burris, PhD and current President Louis J. Muglia, MD, PhD for their support and encouragement to pursue the scholarly analysis of the Career Awards at the Scientific Interface program. This paper was written using data obtained on December 5, 2018, from Digital Science’s Dimensions platform, available at https://app.dimensions.ai. Access was granted to subscription-only data sources under the license agreement.

Additionally, they thank Rusty Kelley, PhD, MBA, former CASI Program Officer, for his initial undertaking of this project and support and review of the manuscript. Finally, the authors thank the CASI applicants and awardees for their dedication and commitment to the advancement of interdisciplinary sciences.

## Supporting Information

**S1 Table. Investigator Distributions by Year and by Cohort.** Breakdown of number of investigators in each category from analyzed in this study by award year.

**S2 Table. Investigator Distribution By Cohort Status and Year.** Breakdown of number of investigators in each category from analyzed in this study by award year 2002-2017.

## Notes

### Competing Interest Statement

The authors have declared no competing interest.

